# Characterization of the SARS-CoV-2 BA.5.5 and BQ.1.1 Omicron Variants in Mice and Hamsters

**DOI:** 10.1101/2023.04.28.538747

**Authors:** James Brett Case, Suzanne M. Scheaffer, Tamarand L. Darling, Traci L. Bricker, Lucas J. Adams, Houda Harastani, Reed Trende, Shilpa Sanapala, Daved H. Fremont, Adrianus C. M. Boon, Michael S. Diamond

**Affiliations:** Department of Medicine,Washington University School of Medicine, St. Louis, MO; Department of Pathology & Immunology, Washington University School of Medicine, St. Louis, MO; Department of Molecular Microbiology,Washington University School of Medicine, St. Louis, MO; Department of Biochemistry and Molecular Biophysics, Washington University School of Medicine, St. Louis, MO; Andrew M. and Jane M. Bursky Center for Human Immunology and Immunotherapy Programs, Washington University School of Medicine, Saint Louis, MO; Center for Vaccines and Immunity to Microbial Pathogens, Washington University School of Medicine, Saint Louis, MO

**Keywords:** SARS-CoV-2, coronavirus, pathogenesis, BA.5.5, BQ.1.1, variants

## Abstract

The continued evolution and emergence of novel SARS-CoV-2 variants has resulted in challenges to vaccine and antibody efficacy. The emergence of each new variant necessitates the need to re-evaluate and refine animal models used for countermeasure testing. Here, we tested a currently circulating SARS-CoV-2 Omicron lineage variant, BQ.1.1, in multiple rodent models including K18-hACE2 transgenic, C57BL/6J, and 129S2 mice, and Syrian golden hamsters. In contrast to a previously dominant BA.5.5 Omicron variant, inoculation of K18-hACE2 mice with BQ.1.1 resulted in a substantial weight loss, a characteristic seen in pre-Omicron variants. BQ.1.1 also replicated to higher levels in the lungs of K18-hACE2 mice and caused greater lung pathology than the BA.5.5 variant. However, C57BL/6J mice, 129S2 mice, and Syrian hamsters inoculated with BQ.1.1 showed no differences in respiratory tract infection or disease compared to animals administered BA.5.5. Airborne or direct contact transmission in hamsters was observed more frequently after BQ.1.1 than BA.5.5 infection. Together, these data suggest that the BQ.1.1 Omicron variant has increased virulence in some rodent species, possibly due to the acquisition of unique spike mutations relative to other Omicron variants.

**IMPORTANCE:** As SARS-CoV-2 continues to evolve, there is a need to rapidly assess the efficacy of vaccines and antiviral therapeutics against newly emergent variants. To do so, the commonly used animal models must also be reevaluated. Here, we determined the pathogenicity of the circulating BQ.1.1 SARS-CoV-2 variant in multiple SARS-CoV-2 animal models including transgenic mice expressing human ACE2, two strains of conventional laboratory mice, and Syrian hamsters. While BQ.1.1 infection resulted in similar levels of viral burden and clinical disease in the conventional laboratory mice tested, increases in lung infection were detected in human ACE2-expressing transgenic mice, which corresponded with greater levels of pro-inflammatory cytokines and lung pathology. Moreover, we observed a trend towards greater animal-to-animal transmission of BQ.1.1 than BA.5.5 in Syrian hamsters. Together, our data highlight important differences in two closely related Omicron SARS-CoV-2 variant strains and provide a foundation for evaluating countermeasures.

## INTRODUCTION

The COVID-19 pandemic caused by severe acute respiratory syndrome coronavirus 2 (SARS-CoV-2) is now in its third year. While major advances have been made in understanding the biology of SARS-CoV-2, the efficacy of existing vaccines and therapeutics has been challenged by the continued evolution of the virus spike glycoprotein. In late 2021, the first Omicron (BA.1) variant emerged, which encoded a greater number of spike mutations than previously detected variants (> 30 mutations relative to WA1/2020). Despite the considerable escape from natural, hybrid, and vaccine-induced immunity in humans, BA.1 caused attenuated infection and disease in mice, hamsters, and non-human primates (1–3). Subsequently, the BA.1 variant split into multiple lineages (BA.2 – 2.75, BA.4, and BA.5), each encoding a unique set of mutations. Since October of 2022, cases of the BQ.1.1 variant, a descendent of the BA.5 lineage, have steadily increased. Relative to BA.5.5, BQ.1.1 encodes only four differences in the spike protein (I76T, R346T, K444T, and N460K) (4).

Here, we evaluated the pathogenesis of the SARS-CoV-2 BQ.1.1 variant relative to the closely related, but genetically distinct, BA.5.5 variant in commercially available mouse strains and Syrian golden hamsters that have been used previously as models of SARS-CoV-2 pathogenesis. For each mouse model, we assessed clinical disease and the levels of viral infection. Using a hamster model, we also evaluated the relative transmissibility of BA.5.5 and BQ.1.1. Finally, we used *in vitro* protein-protein interaction assays to investigate whether the disparities in pathogenesis we observed between variants might be explained by differences in RBD-receptor interactions.

## RESULTS

### BQ.1.1 infection in C57BL/6J, 129S2, and K18-hACE2 mice

The naturally occurring SARS-CoV-2 spike N501Y mutation enables engagement of endogenous murine ACE2 and productive infection of wild-type C57BL/6J and 129S2 mice (5-9). Nonetheless, C57BL/6J, 129S2, and BALB/c mice inoculated with an earlier Omicron variant strain (BA.1) that has the N501Y mutation sustained low levels of viral infection and no clinical disease (1). To determine the pathogenicity of the newly emerged BQ.1.1 variant in these laboratory strains of mice, we inoculated 8-to 10-week-old female C57BL/6J or 16-week-old male 129S2 mice with 10^4^ FFU of BA.5.5 or BQ.1.1. For both C57BL/6J and 129S2 mice, weight loss was not observed after inoculation with BA.5.5 or BQ.1.1 over a four-day period (**Fig 1A-B**), consistent with phenotypes observed with other Omicron strains (1). On day 4 post-infection (dpi), tissues were collected, and the levels of viral infection were determined. In C57BL/6J mice, viral RNA levels in the nasal washes, nasal turbinates, and lungs of animals infected with BA.5.5 or BQ.1.1 were at or barely above the limit of detection, and infectious virus was not recovered from the lungs (**Fig 1A**). Although viral RNA levels in the nasal washes of 129S2 mice were barely detectable (**Fig 1B**), appreciable amounts of BA.5.5 and BQ.1.1 RNA were measured in the lungs and nasal turbinates, with ∽11-fold (*P* < 0.0001) higher amounts of viral RNA in the lung after BA.5.5 infection. Nonetheless, infectious virus recovered from the lungs of 129S2 mice was at or just above the limit of detection.

**Figure 1.**
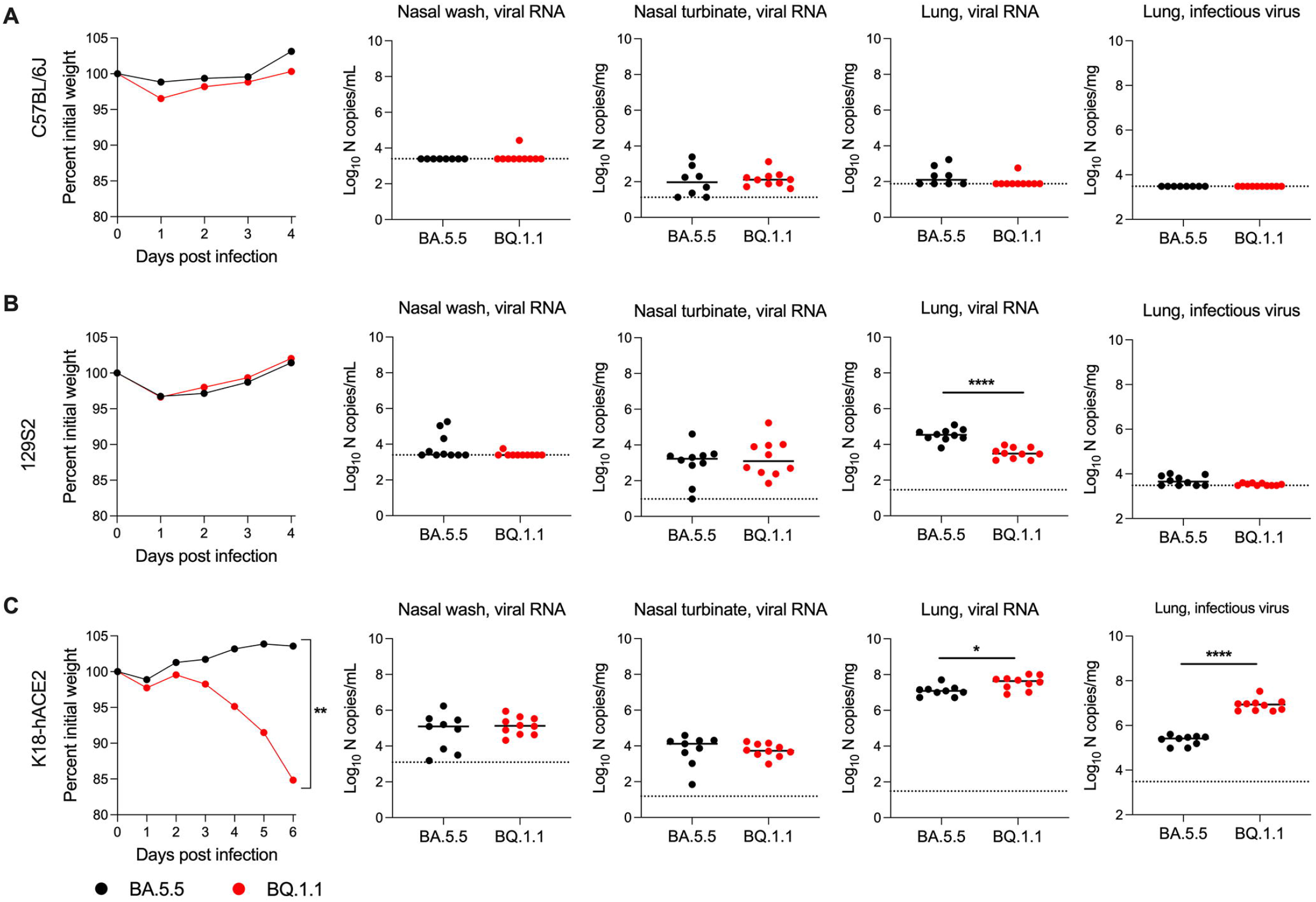
BQ.1.1-infection results in increased infection and weight loss in K18-hACE2 mice. C57BL/6J (**A**), 129S2 (**B**), or K18-hACE2 (**C**) mice were inoculated intranasally with 10^4^ FFU of the indicated SARS-CoV-2 strain. Animals were monitored for weight loss (**A-C**) daily (differences in area under the curves assessed by student’s t-test with Welch’s correction; ** *P* < 0.01). At 4 or 6 dpi, the indicated tissues were collected. Viral RNA levels in the lungs, nasal turbinates, and nasal washes were determined by RT-qPCR, and infectious virus in the lungs were quantified by plaque assay (lines indicate median ± SEM., dotted lines indicate limits of detection; n = 9–10 mice per group, two experiments; Mann-Whitney test between BA.5.5- and BQ.1.1-infected groups; * *P* < 0.05, ****, *P* < 0.0001).

We next tested the capacity of BQ.1.1 and BA.5.5 to infect K18-hACE2 transgenic mice, which are a model of severe disease and pathogenesis for most SARS-CoV-2 strains (10, 11), with the exception of some Omicron isolates (*e.g*., BA.1 and BA.2) that are attenuated (1, 5, 10). After inoculation with 10^4^ FFU, BQ.1.1, but not BA.5.5, resulted in greater than 15% loss in body-weight beginning at 3 dpi (**Fig 1C**). BQ.1.1 infection also resulted in a 3-fold (*P* < 0.05) increase in viral RNA and 33-fold (*P* < 0.0001) increase in infectious virus in the lung compared to the BA.5.5 strain. In the nasal washes and nasal turbinates, comparable levels of viral RNA were observed for both strains. These data suggest the BQ.1.1 variant has an increased capacity to infect the lungs of hACE2-expressing mice, and this correlates with clinical disease as reflected by weight loss.

### BQ.1.1 infection causes increased lung inflammation and pathology in K18-hACE2 mice

Previous reports showed that infection of K18-hACE2 mice with the BA.1 strain did not induce substantive pro-inflammatory cytokine/chemokine responses in the lung (1). Given the capacity of BA.5.5 and BQ.1.1 to replicate in the lungs of K18-hACE2 mice, we quantified the levels of pro-inflammatory cytokines and chemokines present at 6 dpi (**Fig 2A-B**). While increased levels of pro-inflammatory cytokines and chemokines were present in lung homogenates of BA.5.5 or BQ.1.1-infected compared to uninfected mice, many (*e.g*., G-CSF, IL-17, LIF, CCL2, CXCL9, CCL3, and CCL4) were expressed at higher levels in BQ.1.1-than BA.5.5-infected mice. These data suggest that the increased viral replication in the lungs of BQ.1.1-infected K18-hACE2 mice resulted in greater production of pro-inflammatory mediators.

**Figure 2.**
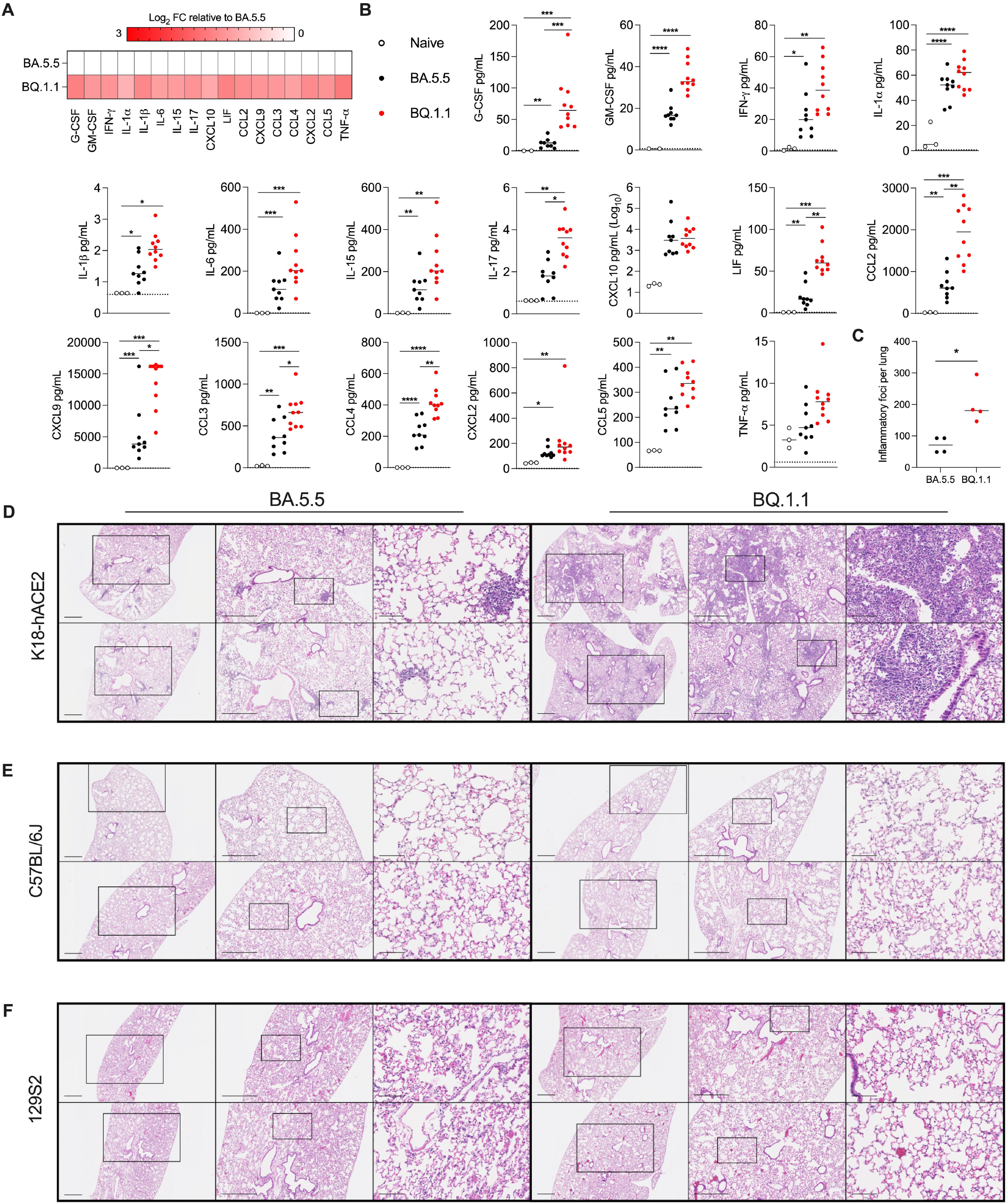
BQ.1.1 infection induces inflammatory cytokines and pathology in the lungs of K18-hACE2 mice. (**A**) Heat map of cytokine and chemokine protein expression levels in lung homogenates. Data from BQ.1.1-infected mice are presented as log_2_-transformed fold-change compared to BA.5.5-infected mice. White, baseline; red, increase. (**B**) Graphs of cytokine and chemokine protein levels in the lungs of naïve, BA.5.5, or BQ.1.1-infected lungs from K18-hACE2 mice at 6 dpi (line indicates median, dotted lines indicate limits of detection; n = 2-3 naïve, n = 9-10 for all other groups (two-way ANOVA with Tukey’s post-test with comparisons between all groups: *, *P* < 0.05, **, *P* < 0.01, ***, *P* < 0.001, ****, *P* < 0.0001). (**C**) Inflammatory foci from mice in **D-F** were quantified blindly and plotted (Mann-Whitney test between BA.5.5- and BQ.1.1-infected groups; * *P* < 0.05). **(D-F**) Hematoxylin and eosin staining of lung sections from K18-hACE2, C57BL/6J, and 129S2 mice collected six days after intranasal inoculation with 10^4^ PFU of the indicated SARS-CoV-2 strain. Images show 2.5× (left), 5× (middle), and 20× (right) power magnification. Scale bars indicate 500 μm, 500 μm, and 100 μm, left-to-right, respectively. Two representative images are shown from four mice per group harvested from two experiments.

To determine the impact of the differences in viral burden and cytokines on lung injury, we performed histopathological analysis of lung tissues using hematoxylin and eosin staining (**Fig 2C-F**). Lungs isolated from K18-hACE2 mice at 6 dpi with BA.5.5 showed limited immune cell infiltration or lung injury. In contrast, BQ.1.1-infected lungs showed evidence of pneumonia with increased numbers of inflammatory lesions characterized by immune cell infiltrates, alveolar space consolidation, vascular congestion, and interstitial edema (**Fig 2C-D**). As expected, lung tissues isolated from C57BL/6J and 129S2 mice infected with BA.5.5 or BQ.1.1 showed no evidence of lung pathology (**Fig 2E-F**).

### BA.5.5. and BQ.1.1 infection and transmission in Syrian hamsters

We evaluated the pathogenicity and transmissibility of BA.5.5 and BQ.1.1 in Syrian hamsters. Animals were inoculated with 2.5 × 10^4^ PFU of BA.5.5 or BQ.1.1, weights were measured daily, and at 3 or 6 dpi, tissues were collected to assess viral infection (**Fig 3A**). Compared to initial weights at the time of inoculation, BA.5.5 and BQ.1.1 did not cause weight loss in hamsters over a 14-day period and had only small, non-significant trends, in weight gain (**Fig 3B**). Viral RNA levels in tissues were generally highest at 3 dpi (**Fig 3C**). Between the two variants, the amounts of viral RNA were comparable with the exception of nasal turbinates at 3 dpi, which showed higher (7.5-fold, *P* < 0.01) BA.5.5 levels than BQ.1.1. In a second analysis of infectious virus by plaque assay, higher (81- and 51-fold, *P* < 0.01) levels of BA.5.5 were detected at 3 dpi in the nasal washes and nasal turbinates but not in the lungs (**Fig 3D**). These data demonstrate that direct inoculation of hamsters with BA5.5 or BQ.1.1 results in similar levels of infection and weight loss, with BA.5.5 exhibiting slightly greater viral burden than BQ.1.1 in the upper respiratory tract.

**Figure 3.**
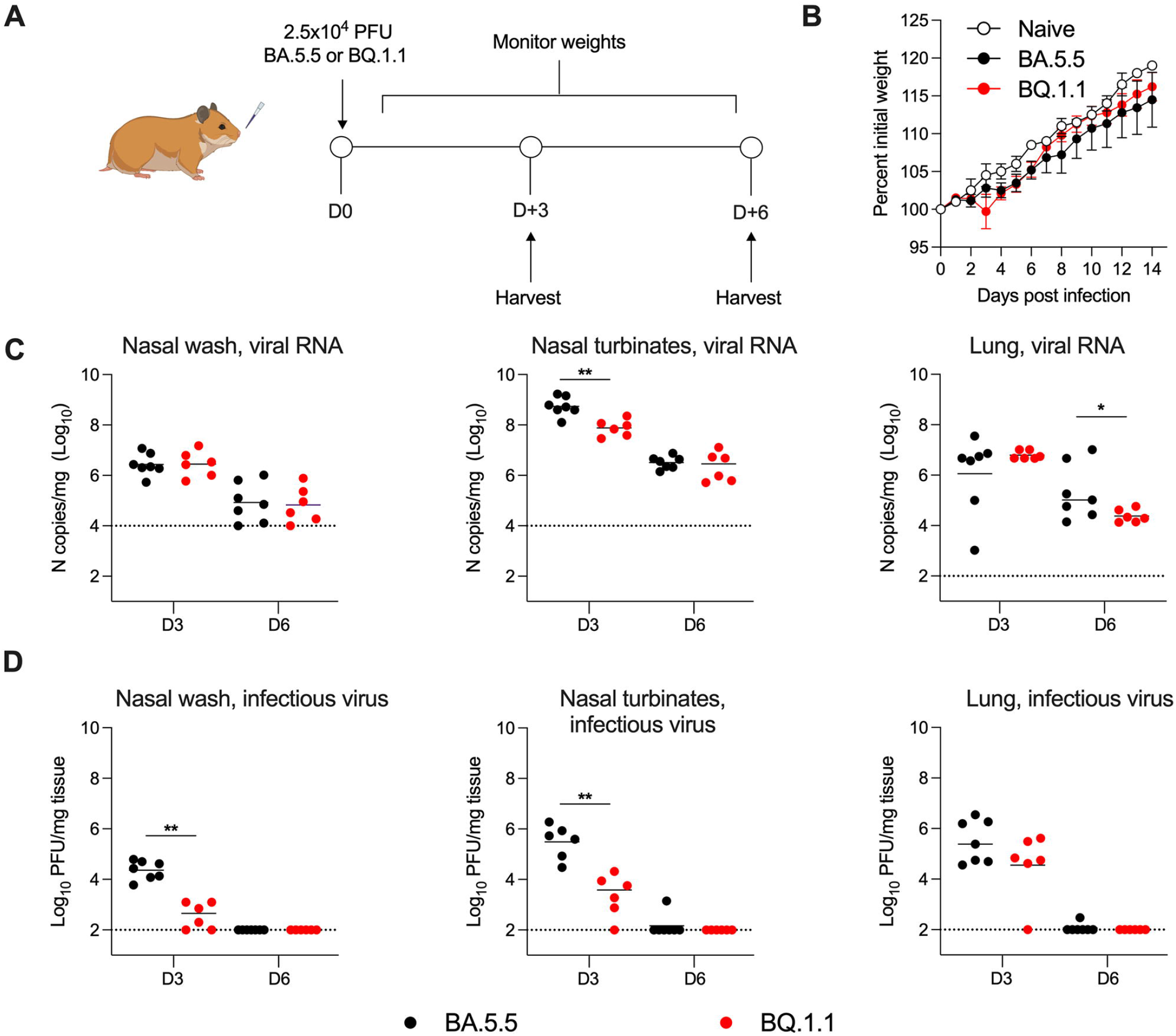
BA.5.5 and BQ.1.1 infections of Syrian hamsters. Hamsters were inoculated intranasally with 2.5 ×10^4^ PFU of the indicated SARS-CoV-2 strain (**A**). Naïve and BA.5.5 or BQ.1.1-inoculated animals were monitored for weight change daily for 14 days (**B**). At 3 or 6 dpi, the nasal washes, nasal turbinates, and lungs, were collected from each animal and levels of viral RNA and infectious virus were determined by RT-qPCR (**C**) or plaque assay (**D**), respectively (lines indicate median ± SEM., dotted lines indicate limits of detection; n = 6-7 hamsters per group per timepoint, two experiments; Mann-Whitney test between BA.5.5- and BQ.1.1-infected groups; * *P* < 0.05, **, *P* < 0.01).

Unlike mice, the SARS-CoV-2 pathogenesis in Syrian hamsters also allows for assessment of viral transmission from animal-to-animal (12, 13). We evaluated the capacity of BA.5.5 and BQ.1.1 to transmit to naïve hamsters via direct contact or airborne transmission (**Fig 4A-B**). Donor hamsters were inoculated with 2.5 ×10^4^ PFU of BA.5.5 or BQ.1.1. At 24 h post-infection, donor animals were transferred to cages containing naïve animals for 8 h of either direct contact or placed inside porous stainless-steel isolation canisters with directional airflow from the donor animal to the naïve animal for airborne transmission. After the 8 h period, donor and contact animals were separated, returned to individual caging, and respiratory tissues were collected 4 days later for detection of infectious virus. Whereas BA.5.5 was transmitted approximately 67% (4 of 6 animals) of the time under direct contact conditions, airborne transmission was not observed (0 of 8 animals) for this variant (**Fig 4C-E**). In contrast, BQ.1.1 was transmitted to recipient animals at frequencies of 100% (6 of 6 animals) and 17% (1 of 6 animals) in direct contact and airborne settings, respectively (**Fig 4C-E**). The difference in direct contact or airborne transmission between the two variants of SARS-CoV-2 was not statistically significant, but trended toward greater transmission of BQ.1.1.

**Figure 4.**
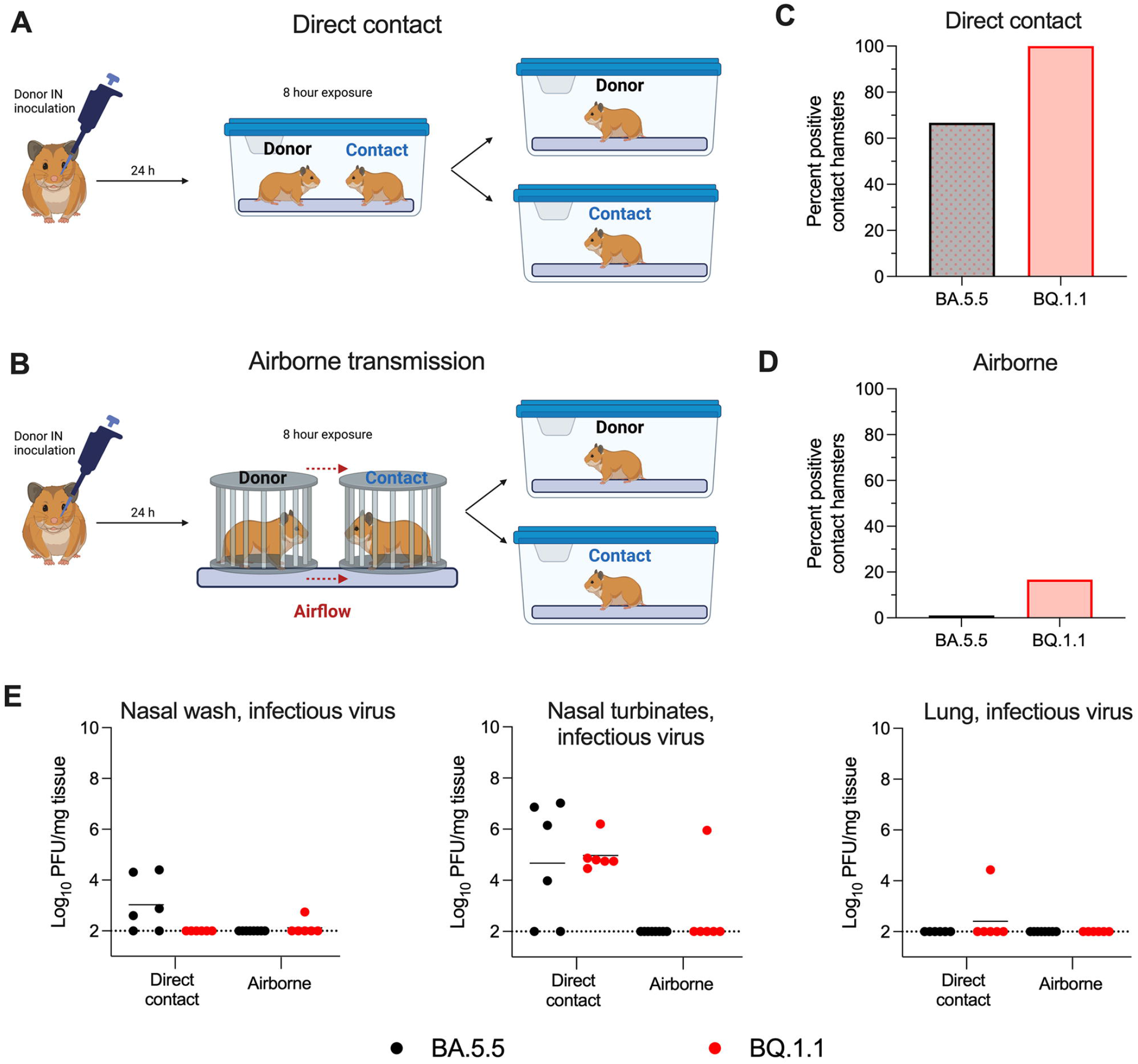
BA.5.5 and BQ.1.1 transmission in Syrian hamsters. For transmission studies, donor hamsters were inoculated intranasally with 2.5 ×10^4^ PFU of the indicated SARS-CoV-2 strain. At 24 h post-inoculation, animals were transferred to cages containing naïve contact animals (direct contact; **A**) or porous canisters (airborne; **B**) upwind of naïve contact animals for a total exposure time of 8 h. After exposure, animals were returned to individual cages. At 4 days post-exposure, the percentage of SARS-CoV-2 positive contact hamsters were quantified (**C** and **D**). Nasal washes, nasal turbinates, and lungs, were collected from contact animals and levels of infectious virus were determined (**E**) (lines indicate median ± SEM., dotted lines indicate limits of detection; n = 6–7 hamsters per group, two experiments). Positive transmission events were registered when infectious virus was detected above the limit of detection within any tissue for a given animal.

### Binding affinities of SARS-CoV-2 BA.5.5 and BQ.1.1 RBDs for human and mouse ACE2

As the SARS-CoV-2 spike protein has evolved in variant strains, differences in binding affinity for hACE2 have been observed (14-16). Since we observed an elevated lung viral burden for BQ.1.1 in K18-hACE2 mice, which express high levels of hACE2 in epithelial cells (10, 17), but not in C57BL/6J or 129S2 mice, which express mouse ACE2 but not hACE2, we performed biolayer interferometry (BLI) experiments to quantify RBD-ACE2 binding interactions (**Fig 5A-C**). For BA.5.5 and BQ.1.1 RBDs, the affinities for hACE2 were similar in the low nanomolar-range (6.06 and 4.4 nM, respectively). In contrast, the affinity of BQ.1.1 for mouse ACE2 (mACE2) was more than 5-fold higher than BA.5.5 (23 versus 121 nM, respectively). Thus, the increased viral burden observed during BQ.1.1 infection in K18-hACE2, but not C57BL/6J or 129S2 mice, is likely not explained by altered affinity of binding to hACE2.

**Figure 5.**
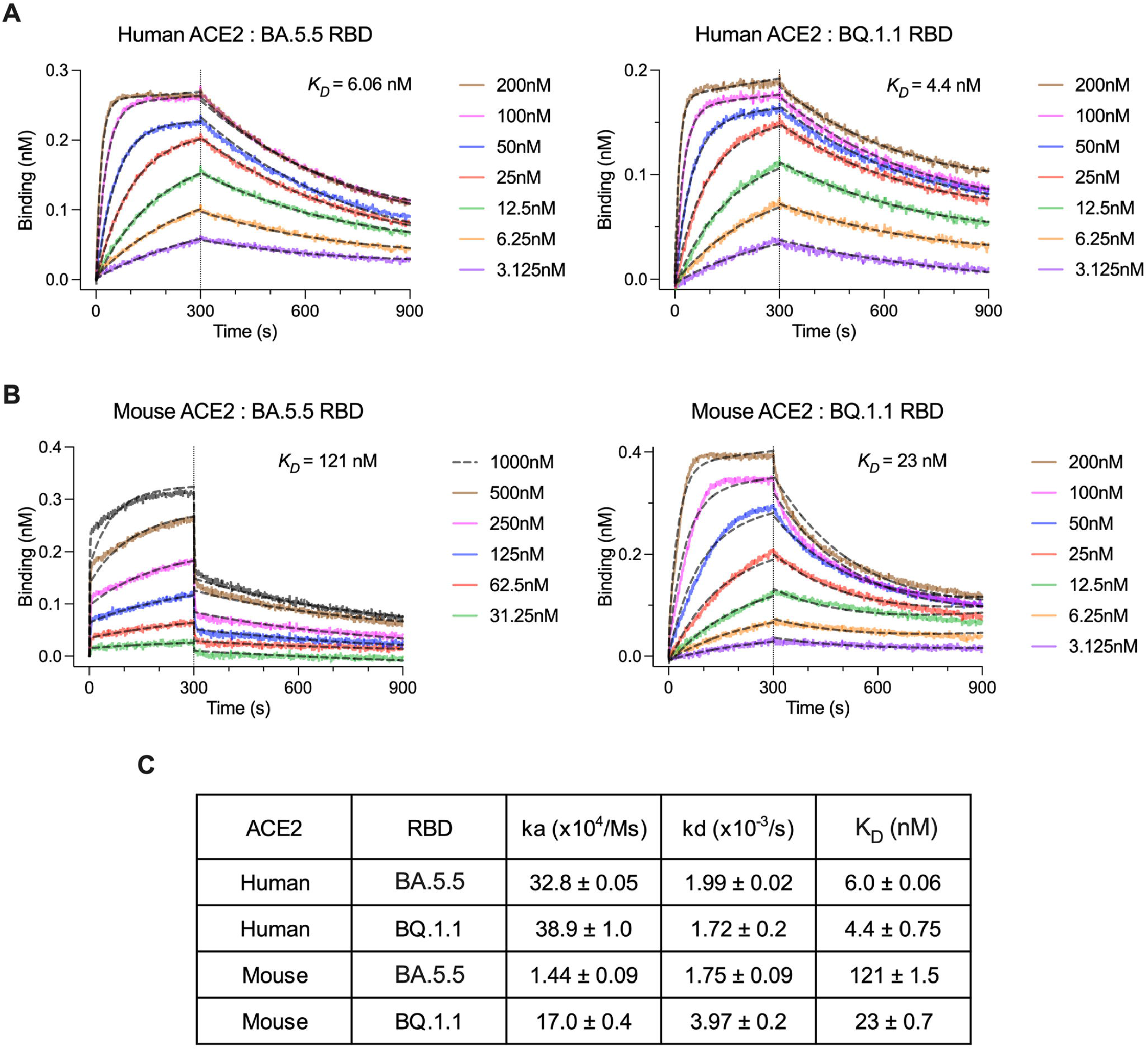
Determination of ACE2-BA.5.5/BQ.1.1 RBD binding affinity by BLI. Recombinant human (**A**) or mouse (**B**) ACE2-Fc proteins were loaded onto biolayer interferometry (BLI) protein G pins at a concentration of 10 μg/mL and dipped into the indicated concentrations of BA.5.5 or BQ.1.1 RBDs. Samples were allowed to associate and dissociate for 300 s and 600 s, respectively. Dashed black curves show fits to a 1:1 binding model with a drifting baseline. (**C**) Association rate (ka), dissociation rate (kd), and kinetic dissociation constant (K_D_) values were calculated and reported.

## DISCUSSION

In this study, we compared the pathogenicity of a circulating SARS-CoV-2 Omicron variant BQ.1.1 to the closely related BA.5.5 variant, the latter of which is rapidly diminishing in prevalence (4). Our experiments highlight phenotypic differences among Omicron variant strains in different animal models commonly used for countermeasure testing. For instance, infection with BA.1 and BA.2 Omicron variant strains results in decreased infection and attenuated disease in mice and hamsters (1, 14). Our studies showed that BQ.1.1 infection of K18-hACE2 transgenic mice resulted in weight loss that was similar to that seen with of pre-Omicron SARS-CoV-2 variants including WA1/2020 D614G, B.1.1.7, and B.1.351 (5, 18), whereas, BA.5.5 infection did not result in weight loss regardless of the model tested. While the exact mechanism of selective BQ.1.1-, but not BA5.5-induced weight loss in K18-hACE2 mice remains to be determined, it may be due in part to the relatively higher levels of infection and inflammation, which correlate with clinical disease progression in K18-hACE2 mice (11).

We observed increased viral burden and pathogenicity in the lungs of BQ.1.1-infected K18-hACE2 mice compared to BA.5.5-infected animals. One potential mechanism that may, in part, explain this observation is differences in mouse and human ACE2 receptor affinity due to the mutations present in BQ.1.1, but not BA.5.5, spike proteins (I76T, R346T, K444T, and N460K). Indeed, a recent report suggested that BQ.1.1 RBD had increased affinity for hACE2 in a yeast surface display assay (1.5-fold), and this correlated with greater cell infectivity and fusogenicity *in vitro* compared to BA.5 (19). In agreement with these data, our BLI results showed that BQ.1.1 RBD had a 1.5-fold greater affinity for hACE2 than BA.5.5, suggesting a mechanism that might contribute to and/or explain the phenotypic differences in K18-hACE2 mice. Reverse genetic experiments with amino acid substituted BA.5 RBDs suggest that the R346T mutation alone is sufficient to enhance affinity for hACE2, whereas R346T and N460K are required to increase *in vitro* cell infectivity (19). Nonetheless, amino acid differences in other structural and nonstructural proteins apart from spike (*e.g*., nsp2: Q376K, nsp6: L260F, nsp12: Y273H, nsp13: M233I, N268S, and N: E136D) also might play a role. This hypothesis that amino acid substitutions outside of spike may be important in distinguishing BA.5.5 and BQ.1.1 infectivity is supported by our observation that a 5-fold increase in BQ.1.1 RBD affinity for mACE2 was insufficient to promote greater infection of C57BL/6J and 129S2 mice compared to BA.5.5. Furthermore, the observation of a trend toward increased transmissibility for BQ.1.1 relative to BA.5.5 in hamsters suggests that more studies in animal models and humans are needed to fully discern differences in pathogenesis for these and other SARS-CoV-2 variants.

### Limitations of study

Several limitations exist in our study: (1) We evaluated three mouse strains and a single Syrian hamster model of SARS-CoV-2 disease. Although these animal models are used extensively in studying SARS-CoV-2 pathogenesis and countermeasures, future testing of the virulence of BQ.1.1 and BA.5.5 in non-human primates will be important; (2) While the hamster transmission model is useful, it will be important to corroborate our BQ.1.1 transmission findings in humans as data becomes available; (3) All of our studies were performed in naïve animals, which does not address the relative immune evasive potential of BQ.1.1 and BA.5.5, a key question given that most of the global population has been infected or immunized; and (4) We did not evaluate even newer strains of the XBB.1 lineage. During the latter stages of our study, cases of SARS-CoV-2 associated with the XBB lineage have increased substantially. Future studies are needed to compare the pathogenicity, transmissibility, and immune evasive potential of XBB and BQ.1.1 strains in animal models and humans.

## ACKNOWLEDGEMENTS

This study was supported by the NIH (R01 AI157155, NIAID Centers of Excellence for Influenza Research and Response (CEIRR) contracts 75N93021C00014 and 75N93019C00051, to M.S.D., 75N93021C00016 to A.C.M.B, and 75N93022C00035 and 75N93019C00062 to D.H.F. J.B.C is supported by a Moderna Global Fellowship award. We thank Aaron Schmidt and Catherine Jacob-Dolan for sharing the BQ.1.1 RBD protein used for BLI studies, and Mehul Suthar and Andrew Pekosz for BA.5.5 and BQ.1.1 isolates, respectively. Some figure designs were created using BioRender.com.

## AUTHOR CONTRIBUTIONS

J.B.C. and S.M.S. performed mouse experiments and viral burden analyses. S.M.S. propagated and validated SARS-CoV-2 viruses. T.L.D., T.L.B., H.H., and R.T. performed hamster experiments. T.L.D., T.L.B., and H.H. performed hamster viral burden analyses. L.J.A., J.B.C., and S.S. performed BLI experiments. S.S. performed histopathology scoring. D.H.F., A.C.M.B, and M.S.D. obtained funding and supervised the research. J.B.C. and M.S.D. wrote the initial draft, with the other authors providing editorial comments.

## COMPETING FINANCIAL INTERESTS

M.S.D. is a consultant for Inbios, Vir Biotechnology, Ocugen, Topspin, Moderna, and Immunome. The Diamond laboratory has received unrelated funding support in sponsored research agreements from Vir Biotechnology, Emergent BioSolutions, Generate Biomedicines, and Moderna. The Boon laboratory has received unrelated funding support in sponsored research agreements from GreenLight Biosciences Inc., Moderna, and AbbVie Inc. All other authors declare no competing financial interests.

## METHODS

### Cells

Vero-TMPRSS2 (20) and Vero-hACE2-TMPRRS2 (21) cells were cultured at 37°C in Dulbecco’s Modified Eagle medium (DMEM) supplemented with 10% fetal bovine serum (FBS), 10□mM HEPES pH 7.3, 1□mM sodium pyruvate, 1× non-essential amino acids, and 100□U/ml of penicillin–streptomycin. Vero-TMPRSS2 cells were supplemented with 5 μg/mL of blasticidin. Vero-hACE2-TMPRSS2 cells were supplemented with 10 μg/mL of puromycin. All cells routinely tested negative for mycoplasma using a PCR-based assay.

### Viruses

The BA.5.5 (hCoV-19/USA/COR-22-063113/2022) and BQ.1.1 strains (hCoV-19/USA/CA-Stanford-79_S31/2022) were obtained from nasopharyngeal isolates as generous gifts of Andrew Pekosz (Johns Hopkins School of Public Health) and Mehul Suthar (Emory University), respectively. All virus stocks were generated in Vero-TMPRSS2 cells and subjected to next-generation sequencing as described previously (21) to confirm the presence and stability of expected substitutions. All virus experiments were performed in an approved biosafety level 3 (BSL-3) facility.

### Mouse experiments

Animal studies were carried out in accordance with the recommendations in the Guide for the Care and Use of Laboratory Animals of the National Institutes of Health. The protocols were approved by the Institutional Animal Care and Use Committee at the Washington University School of Medicine (assurance number A3381–01). Virus inoculations were performed under anesthesia that was induced and maintained with ketamine hydrochloride and xylazine, and all efforts were made to minimize animal suffering.

Heterozygous K18-hACE2 C57BL/6J mice (strain: 2B6.Cg-Tg(K18-ACE2)2Prlmn/J), and wild-type C57BL/6J (strain: 000664) were obtained from The Jackson Laboratory. 129S2 mice (strain: 129S2/SvPasCrl) were obtained from Charles River Laboratories. All animals were housed in groups and fed standard chow diets. For mouse experiments, eight-to ten-week-old female K18-hACE2 and C57BL/6J mice or 16-week-old male 129S2 mice were administered the indicated doses of the respective SARS-CoV-2 strains by intranasal administration. *In vivo* studies were not blinded, and mice were randomly assigned to treatment groups. No sample-size calculations were performed to power each study. Instead, sample sizes were determined based on prior *in vivo* virus challenge experiments.

### Measurement of viral RNA levels

Tissues were weighed and homogenized with zirconia beads in a MagNA Lyser instrument (Roche Life Science) in 1 mL of DMEM medium supplemented with 2% heat-inactivated FBS. Tissue homogenates were clarified by centrifugation at 10,000 rpm for 5 min and stored at −80°C. RNA was extracted using the MagMax mirVana Total RNA isolation kit (Thermo Fisher Scientific) on the Kingfisher Flex extraction robot (Thermo Fisher Scientific). RNA was reverse transcribed and amplified using the TaqMan RNA-to-CT 1-Step Kit (Thermo Fisher Scientific). Reverse transcription was carried out at 48°C for 15 min followed by 2 min at 95°C. Amplification was accomplished over 50 cycles as follows: 95°C for 15 s and 60°C for 1 min. Copies of SARS-CoV-2 *N* gene RNA in samples were determined using a previously published assay (22). Briefly, a TaqMan assay was designed to target a highly conserved region of the *N* gene (Forward primer: ATGCTGCAATCGTGCTACAA; Reverse primer: GACTGCCGCCTCTGCTC; Probe: /56-FAM/TCAAGGAAC/ZEN/AACATTGCCAA/3IABkFQ/). This region was included in an RNA standard to allow for copy number determination down to 10 copies per reaction. The reaction mixture contained final concentrations of primers and probe of 500 and 100 nM, respectively.

### Viral plaque assay

Vero-TMPRSS2-hACE2 cells were seeded at a density of 1×10^5^ cells per well in 24-well tissue culture plates. The following day, medium was removed and replaced with 200 μL of material to be titrated diluted serially in DMEM supplemented with 2% FBS. One hour later, 1 mL of methylcellulose overlay was added. Plates were incubated for 72 h, then fixed with 4% paraformaldehyde (final concentration) in PBS for 20 min. Plates were stained with 0.05% (w/v) crystal violet in 20% methanol and washed twice with distilled, deionized water.

### Hamster experiments

Five-six-week-old male hamsters were obtained from Charles River Laboratories and directly transferred into an enhanced animal biosafety level 3 laboratory (ABSL-3+). Hamsters were challenged via the intranasal route with 2.5 ×10^4^ PFU of SARS-CoV-2 BA.5.5 or BQ.1.1. Twenty-four hours after challenge, naïve contact hamsters were exposed to the directly inoculated (donor) hamsters. Potential virus exposure was either airborne, wherein the donor and contact hamsters were placed in individual porous stainless-steel isolation canisters with directional airflow coming from the donor cage, or through direct contact, wherein the contact hamster was placed directly into the donor cage. Upon eight hours of exposure, the contact hamsters were returned to their original cages. Hamsters directly inoculated were weighed daily and harvested at three- or six-days post-inoculation. Contact hamsters were harvested at four days post exposure. At time of harvest, the left lung lobe, nasal wash, and nasal turbinate of each animal was collected. The nasal wash was collected with 1 mL of PBS supplemented to contain 0.1% BSA and subsequently clarified at 1,200 rpm for ten minutes at 4°C. The left lung lobe was homogenized in 1.0 mL DMEM and clarified by centrifugation 1,000 × g for 5 min. Nasal turbinates were collected by removing the skin along the nose and cheeks followed by cutting the jaw to expose the upper palate. A sagittal incision through the palate exposed the nasal turbinates which were then removed using blunt forceps. The nasal turbinates were homogenized in 1.0 mL of DMEM supplemented with 2% FBS, 10mM HEPES (pH 7.3) and 2 mM L-glutamine and clarified by centrifugation 1,000 × g for 5 min. The nasal wash, lung and nasal turbinates were used for viral titer analysis by quantitative RT-PCR using primers and probes targeting the N gene, and by plaque assay. Infectious viral titer detected in any of the contact hamster tissues was considered a positive transmission event.

### Cytokine and chemokine protein measurements

Lung homogenates were incubated with Triton-X-100 (1% final concentration) for 1 h at room temperature to inactivate SARS-CoV-2. Homogenates were analyzed for cytokines and chemokines by Eve Technologies Corporation (Calgary, AB, Canada) using their Mouse Cytokine Array/Chemokine Array platform.

### Lung pathology

Animals were euthanized before harvest and fixation of tissues. Briefly, lungs were inflated with approximately 1.2 mL of 4% paraformaldehyde using a 3-mL syringe and catheter inserted into the trachea. Tissues were allowed to fix for 24 h at room temperature, embedded in paraffin, and sections were stained with hematoxylin and eosin. Slides were scanned using a Hamamatsu NanoZoomer slide scanning system, and the images were viewed using NDP view software (ver.1.2.46). Inflammatory lesions were quantified blindly.

### Binding analysis by biolayer interferometry

BLI was used to quantify the binding of BA.5.5 and BQ.1.1 SARS-CoV-2 RBDs to recombinant human or mouse ACE2 proteins. Fc conjugated human and mouse ACE2 proteins were expressed and purified as previously described (23). Subsequently, 10 μg/mL of each protein was immobilized onto protein G biosensors (GatorBio) for 3 min. After a 30 s wash, the pins were submerged in running buffer (10 mM HEPES, 150 mM NaCl, 3 mM EDTA, 0.05% P20 surfactant, and 1% BSA) containing BA.5.5 or BQ.1.1 RBD protein (Sino Biologics) ranging from 3.125 to 1,000 nM, followed by a dissociation step in running buffer alone. The BLI signal was recorded and analyzed as a 1:1 binding model with a drifting baseline using BIAevaluation Software (Biacore).

### Data availability

All data supporting the findings of this study are available within the paper and are available from the corresponding author upon request.

### Statistical analysis

All statistical tests were performed as described in the indicated figure legends using Prism 9.4.1. Statistical significance was determined using an ANOVA when comparing three or more groups. When comparing two groups, a Mann-Whitney test was performed. The number of independent experiments performed are indicated in the relevant figure legends.

